# Assessment of Brain Glucose Metabolism Following Cardiac Arrest by [^18^F]FDG Positron Emission Tomography

**DOI:** 10.1101/2020.01.08.899252

**Authors:** Hannah J. Zhang, Samuel Mitchell, Yong-Hu Fang, Hsiu-Ming Tsai, Lin Piao, Alaa Ousta, Lara Leoni, Chin-Tu Chen, Willard W. Sharp

**Author notes:** **Corresponding author** Willard W. Sharp PhD, MD; Department of Medicine, Section of Emergency Medicine, University of Chicago, 5841 S Maryland Avenue, Chicago, IL 60637. The Work was done at the University of Chicago.

## Abstract

**Background:** Cardiac arrest (CA) patients who survived by cardiopulmonary resuscitation (CPR) can present different levels of neurological deficits ranging from minor cognitive impairments to persistent vegetative state and brain death. The pathophysiology of the resulting brain injury is poorly understood and whether changes in post-CA brain metabolism contribute to the injury are unknown. Here we utilized [^18^F]FDG-PET to study *in vivo* cerebral glucose metabolism 72 hours following CA in a murine cardiac arrest model.

**Methods:** Anesthetized and ventilated adult C57BL/6 mice underwent 12-minute KCl-induced CA followed by CPR. Seventy-two hours following cardiac arrest, surviving mice were intraperitoneally injected with [^18^F]FDG (~186 μCi/200 μL) and imaged on Molecubes preclinical micro PET/CT imaging systems after a 30-minute awake uptake period. Brain [^18^F]FDG uptake was determined by the VivoQuant software on fused PET/CT images with the 3D brain atlas. Upon completion of PET imaging, remaining [^18^F]FDG radioactivity in the brain, heart, and liver was determined using a gamma counter.

**Results:** Global increases in brain [^18^F]FDG uptake in post-CA mice were observed compared to shams and controls. The median standardized uptake value (SUV) of [^18^F]FDG for CA animals was 1.79 vs. sham 1.25 (p<0.05) and control animals 0.78 (p<0.01). This increased uptake was consistent throughout the 60-minute imaging period and across all brain regions reaching statistical significance in the midbrain, pons, and medulla. Biodistribution analyses of various key organs yielded similar observations that the median [^18^F]FDG uptake for brain were 7.04%ID/g tissue for CA mice vs 5.537%ID/g tissue for sham animals, p<0.05).

**Conclusions:** This study has successfully applied [^18^F]FDG-PET/CT to measure changes in brain metabolism in a murine model of asystolic CA. Our results demonstrate increased [^18^F]FDG uptake in the brain 72 hours following CA, suggesting increased metabolic demand in the case of severe neurological injury. Further study is warranted to determine the etiology of these changes.

## Introduction

There are approximately 550,000 cardiac arrests (CA) both in and out of the hospital each year in the United States^1^. Despite improvements in cardiopulmonary resuscitation, morbidity and mortality in successfully resuscitated patients is high^2,3^. Although multi-organ injury is not uncommon, many post-CA patients die primarily in the context of severe hypoxic encephalopathy^4^. Currently post-cardiac arrest hypothermia and supportive care are the only treatments for managing post-CA neurological injury. The pathophysiology of post-cardiac arrest neurological injury is not well understood and there are few studies investigating change in brain metabolism following post-CA resuscitation.

As a glucose analogue, [^18^F]fluorodeoxyglucose ([^18^F]FDG) is an FDA approved positron emission tomography (PET) radiotracer, routinely used in the clinical prognosis and diagnosis of cancer and the determination of overall tissue metabolic demand. Similar to glucose metabolism, once [^18^F]FDG is transported into cells by glucose transporters, it is irreversibly phosphorylated and trapped intracellularly. However, in contrast to glucose, [^18^F]FDG does not undergo further glycolysis due to its lack of a 2-hydroxyl group. Therefore, [^18^F]FDG is not only used in tumor imaging, but is also utilized to evaluate the function of tissues with high glycolytic rates including brains.

To date, there have been only a few pre-clinical investigations examining post-cardiac arrest brain metabolism using [^18^F]FDG-PET in dogs and rats in the first several hours following resuscitation^5,6^. Neither of these studies examined time points distant from the cardiac arrest. Clinically the appearance of ischemic brain injury on imaging studies such as CT often does not occur until several days after the event, depending on the severity of the ischemia. Therefore, it is important to determine glucose metabolism during this late phase when brain injury is beginning to become visible on CT. In addition, neither of these prior studies reported the neurological outcomes of the animals studied, making their observation of alterations in glucose metabolism difficult to associate with observed changes in neurological behavior. In this study we assessed global and regional changes in brain glucose metabolism 72 hours following CA using [^18^F]FDG-PET in a mouse model and reported how these changes relate to the neurological outcomes in the animals studied.

## Material and Methods

### Animals

Mature female C57BL/6 mice with median age of 3.5 months (n=25) were used in the experiments. Animals were housed in The University of Chicago Animal Research Resources Center. The Institutional Animal Care and Use Committee of the University of Chicago, in accordance with National Institutes of Health guidelines, approved all animal procedures. Animals were maintained at 22–24°C on a 12:12-h light-dark cycle and provided food (standard mouse chow) and water ad libitum.

### Animal Models of Cardiac Arrest

In this study, age and weight matched mice were divided into control (n=3), sham (n=8) and CA (n=14) groups. None of the procedures were done to the control animals. Sham animals underwent surgery, anesthesiology and ventilation, in a manner similar to CA animals but did not undergo CA as outlined in the scheme of Figure 1A.

**Figure 1.**
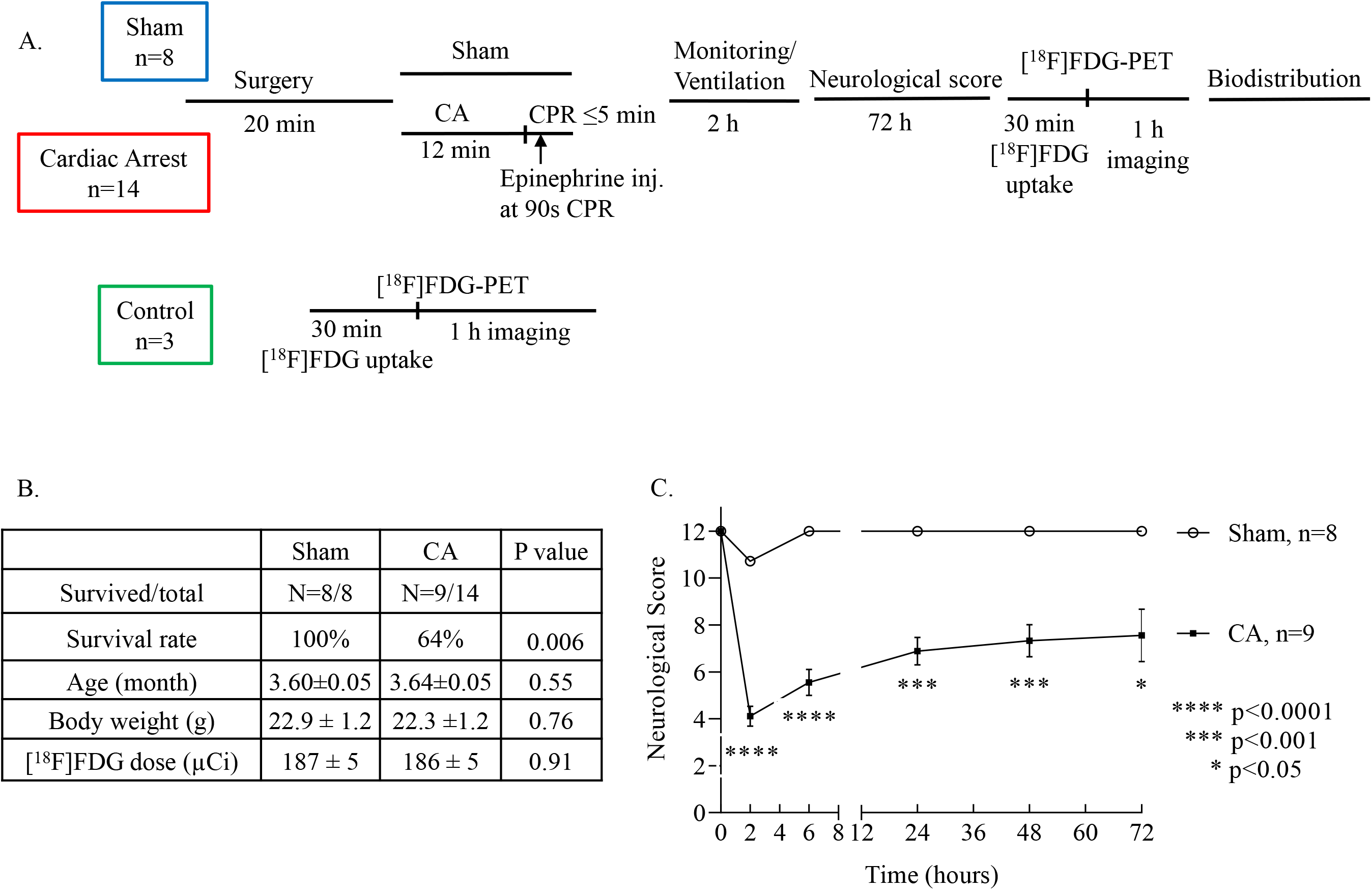
Cardiac arrest induces sustained neuronal damage in resuscitated mice. A) A study time line of animals underwent cardiac arrest (CA) induction and sham treatment. Control animals did not undergo any surgery. B) Table showing animal baseline characteristics and post cardiac arrest survival. Data are presented in mean±SEM. C) Neurological outcome was evaluated according to six preset scoring criteria at 2, 6, 24, 48, and 72 hours following CA/resuscitation. CA animals had a significantly lower average score suggesting an increased neurological damage (mean±SEM).

A modified version of our previously published murine model of asystolic CA was used^7,8^. Briefly, mice were anesthetized with 3% isoflurane and vascular access was acquired. Temperature, respirations, and EKG tracings were recorded continuously on a PowerLab data acquisition module (AD Instruments, Colorado Springs, CO). A 0.4 mm OD heparinized micro PE cannula (BioTime Inc., Berkeley, CA) was placed in the left jugular vein for fluid administration and right carotid artery for aortic systolic pressure (ASP) measurement. Asystolic CA was induced with a single bolus of KCl (0.8 mg/g body weight) into the internal jugular vein and ventilation was suspended. Following 12 min of CA, cardiopulmonary resuscitation (CPR) was performed at approximately 300-350 beats/min. After 90 seconds of CPR, 1.5 μg of epinephrine was injected followed by a saline flush. CPR was terminated when return of spontaneous circulation (ROSC), defined by a mean arterial pressure >40 mm Hg lasting longer than 5 minutes, was achieved. CPR was terminated if ROSC was not achieved after 5 minutes. Quality of CPR was retrospectively evaluated for each animal by reviewing CPR rate and arterial pressure. Resuscitated animals received IV 0.9% saline at a rate of 100 μL/h and were monitored on mechanical ventilation for 120 minutes. Thereafter, the mice were allowed free access to food and water. The mice were observed every 2 hours during the first 6 hours following CA. Animals were then returned to the animal facility.

### Neurological Scoring

Neurological score of mice after CA (2h, 6h, 24h, 48h and 72h) were determined using a 12-point mouse neurological scoring system^7^. Scores ranged from 0 (no response or worst) to 2 (normal) along 6 domains: paw pinch, righting reflex, breathing, spontaneous movement, motor-global and motor-focal. The scores for each of the 6 domains were determined in a blinded fashion and summed to calculate the neurological score.

### [^18^F]FDG-PET/CT Imaging

Seventy-two hours after the surgery, six out of eight surviving sham animals were imaged as well as all of the 9 surviving CA animals in a manner as the control animals (n=3). Two of the sham animals were not imaged due to technical reasons on the day of the experiment. The imaging protocol was designed base on publications from other investigators^9–11^ and our preliminary experiments trialing the protocol (data not shown). Following an overnight fast, all mice (22.6±0.8 g) received 186±3 μCi of [^18^F]FDG (Sofie Biosciences Inc, Dulles, VA) intraperitoneally in 200 μL isotonic saline solution. After a 30-min awake period to facilitate the radiotracer uptake, total body PET imaging was acquired on the β-cube preclinical microPET imaging system (Molecubes, Gent, Belgium) with 133 mm × 72 mm field of view (FOV) and an average spatial resolution of 1.1 mm at the center of FOV^12^. List-mode data was recorded for one hour followed by a reference CT image on the X-cube preclinical microCT imaging system, (Molecubes, Gent, Belgium). The images were reconstructed using an OSEM reconstruction algorithm with an isotropic voxel size of 400 μm and a frame rate of 6×10 minutes. CT images were reconstructed with a 200 μm isotropic voxel size and used for anatomic co-registration and scatter correction. Animals were maintained under 1-2% isoflurane anesthesia in oxygen during imaging. Respiration and temperature were constantly monitored and maintained using the Molecubes monitoring interface, and Small Animal Instruments (SAII Inc, Stoney Brook, NY) set up. All of the animals survived the imaging.

### Data Analyses of [^18^F]FDG Uptake

To quantify [^18^F]FDG uptake, analyses of CT-fused PET images were performed using VivoQuant software (Invicro, Boston, MA). Dynamic [^18^F]FDG uptake was acquired and expressed in standardized uptake value (SUV) for the six 10-minute frames^13,14^

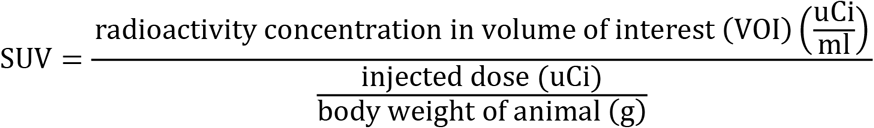

Brain regional analysis was done using a 3-dimensional mouse brain atlas available as the VivoQuant software plug-in. The 3D brain atlas is based on the Paxinos-Franklin atlas registered to a series of high-resolution magnetic resonance imaging with 100 μm near isotropic data and has been applied in other studies^15–17^. The brain was segmented into 14 regions within the skull of each animal using the CT as the reference for automatic registration. The 14 brain regions were the cerebellum, cortex, corpus callosum, hippocampus, hypothalamus, midbrain, medulla, olfactory, pallidum, pons, striatum, thalamus, ventricles, and white matter. The volume of interest (VOI) of the whole brain was 430±17 mm^3^ (mean±SEM). The regional differences of [^18^F]FDG uptake between the CA and the sham were calculated using the following equation.

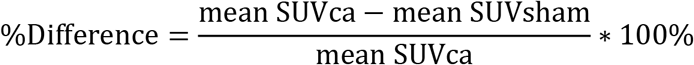

[^18^F]FDG uptake of the heart and the liver was done similarly by drawing the VOI manually with CT guided anatomy. For liver analyses, a 50 mm^3^ spherical VOI was drawn within the boundaries of the right lobe as observed using axial, coronal, and sagittal views of the CT image. For the heart, a spherical VOI with a diameter of 12 pixels was drawn to circumscribe the heart based on the CT image, then a threshold was applied to the PET signal to ensure that VOI only included the myocardium and excluded cardiac chambers. These steps resulted in the VOI of 193±19 mm^3^ (mean±SEM) for the heart.

The relative [^18^F]FDG uptake in the brain was presented as the SUV ratio by dividing the SUV of the whole brain or each brain region to the SUV of the liver of corresponding mouse^18^. In tumor diagnostic imaging, normal liver is often used as the reference region to calculate SUV ratio largely because the perfusion in liver is high^19^. It has been shown that [^18^F]FDG concentration in liver positively correlates with [^18^F]FDG concentration in blood and with the levels of blood glucose^20^.

### Biodistribution of [^18^F]FDG

Biodistribution of [^18^F]FDG uptake was performed on animals after each PET/CT imaging except the animals from the first imaging day. The brains, hearts, and livers from sham (n=5) and CA (n=7) were isolated and weighed. Radioactivity of the organs was measured on a Cobra II gamma counter (PerkinElmer, Waltham, MA). The percentage of uptake (%ID/g tissue) was calculated by dividing the decay-corrected activity in an organ by the total injected dose and normalizing to the weight of the organ.

### Statistical Analyses

Statistical analyses were performed using Prism software (GraphPad, La Jolla, CA, USA). The survival rate between sham and CA animals was evaluated by the Log-rank (Mantel-Cox) test. Continuous data were summarized as media with interquartile range unless it is otherwise indicated specifically, with statistically significant differences assessed using a non-parametric Mann-Whitney test between two groups and Kruskal-Wallis text among multiple groups. The differences of neurological scores between sham and CA groups (means ±SEM) were determined by two-way ANOVA using the Bonferroni post hoc test. The correlation analyses were done using simple linear regression. p<0.05 was considered statistically significant.

## Results

There were no difference in regard to animals’ age, body weight and dose of [^18^F]FDG received (Fig 1B). However, 72 hours after CA the survival rates between sham and CA groups were significantly different (p=0.006). Approximately two third of the CA mice (9 out of 14) survived, compared to survival of all sham animals (8 out of 8). CA mice that survived demonstrated neurological disability as early as 2 hours after resuscitation. At 72 hours, neurological scores in CA mice had improved compared with the scores at 2 hours post CA, but scores were still significantly lower than those of sham mice at the same time point (P<0.05, Figure 1C).

Figure 2A shows the [^18^F]FDG uptake of 5 representative coronal sections from anterior to posterior portion of the brain in sham (bottom) and CA (top) mice. The image at the bottom of coronal sections is a schematic diagram of sagittal view of the brain with 5 perpendicular lines showing the approximate position where the coronal sections were taken. The [^18^F]FDG uptake was consistent over the 1 hour period of the imaging as analyzed by the dynamic reconstruction (Supplemental Fig 1). Therefore, the averaged SUVs from those of six 10-min time frames were used for the comparison of [^18^F]FDG uptake among control, sham and CA mice. On average, the global brain [^18^F]FDG uptake was significantly higher in CA mice than in sham animals, p<0.05 and controls, p<0.01 (Fig 2B). The [^18^F]FDG uptake of liver and heart was also measured simultaneously but no differences were found (Fig 2D & 2F). These observations were confirmed by the ex-vivo biodistribution analyses measuring the radioactivity of individual organs isolated immediately after the imaging (Fig 2C 2E, & 2G).

**Figure 2.**
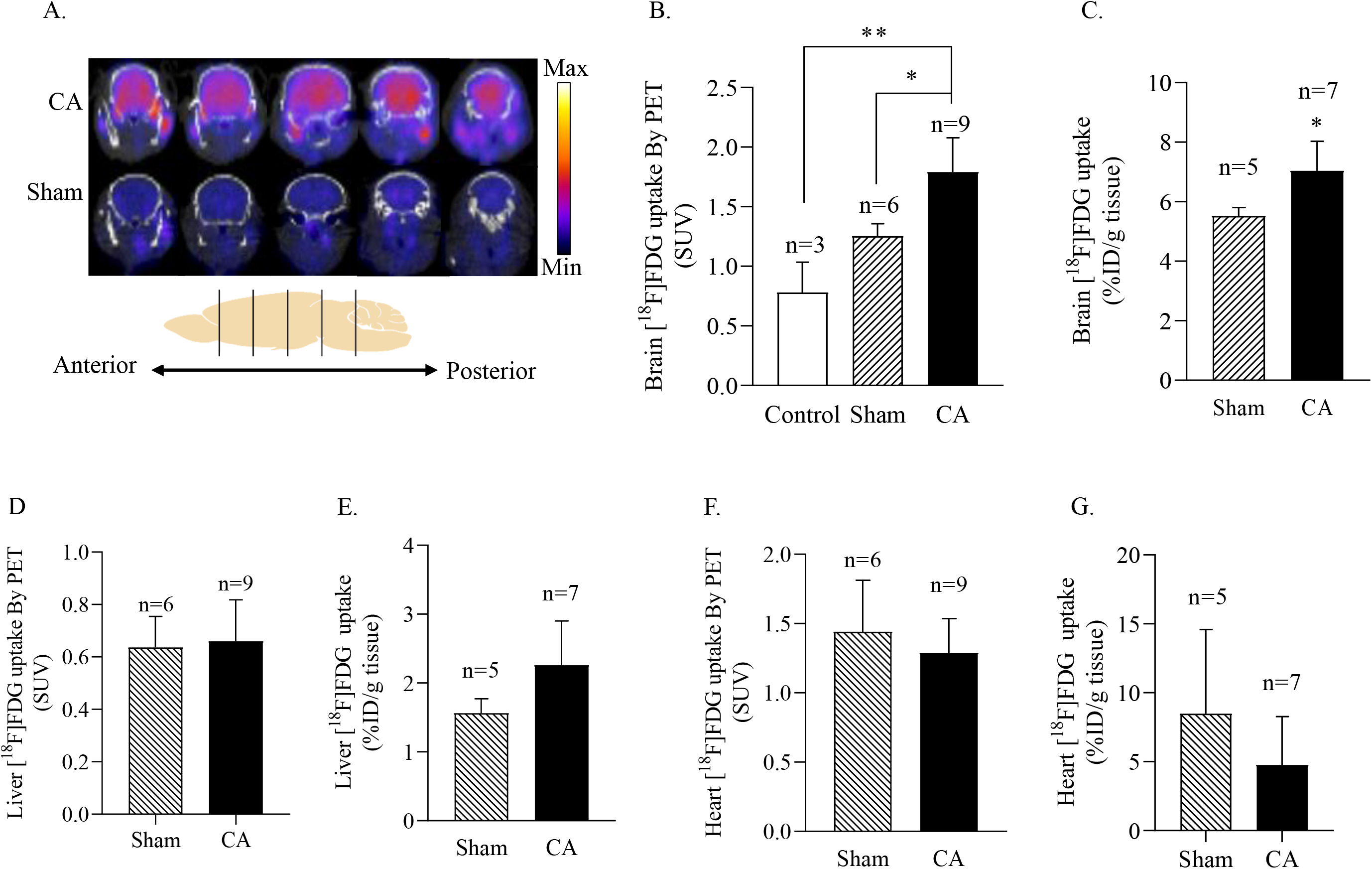
Cardiac arrest increases global brain [^18^F]FDG uptake in mice 72 hours post resuscitation. A) Representative coronal sections from anterior to posterior portions of [^18^F]FDG-PET/CT brain imaging for CA and sham mice. B) Quantification of whole brain [^18^F]FDG uptake as measured by standardized uptake value (SUV) demonstrates a 1.43 and 2.29 fold higher uptake in CA animals vs. sham (p<0.05) and control mice (p<0.01) respectively. C) Ex-vivo average whole brain [^18^F]FDG activity of CA and sham mice as measured by biodistribution (7.04 vs 5.52%ID/g tissue, p<0.05). D) & E) Quantification of liver and heart [^18^F]FDG uptake as measured by SUV. No differences between groups were found. F) & G) Ex-vivo liver and heart [^18^F]FDG activity of CA and sham mice as measured by biodistribution. No differences between groups were found.

To further investigate whether the cerebral [^18^F]FDG uptake was related to the neurological injury demonstrated by the animals, correlation between the neurological scores and the SUV was analyzed by simple linear regression method (Figure 3). It was found that the lower neurological scores were correlated with the higher brain SUV with a statistically significant correlation at the 24 hour (r^2^=0.2722, p=0.0461) (Figure 3A) and 48 hour (r^2^=0.2972, p=0.0356) (Figure 3B) post-CA. A similar trend of correlation was also found for the time points of 2 hours (r^2^=0.0945, p=0.2467) (Supplemental Figure 2A), 6 hours (r^2^=0.1926, p=0.1018) (Supplemental Figure 2B), and 72 hours (r^2^=0.1968, p=0.0977) (Figure 3C) after the CA, but statistical significance was not achieved.

**Figure 3.**
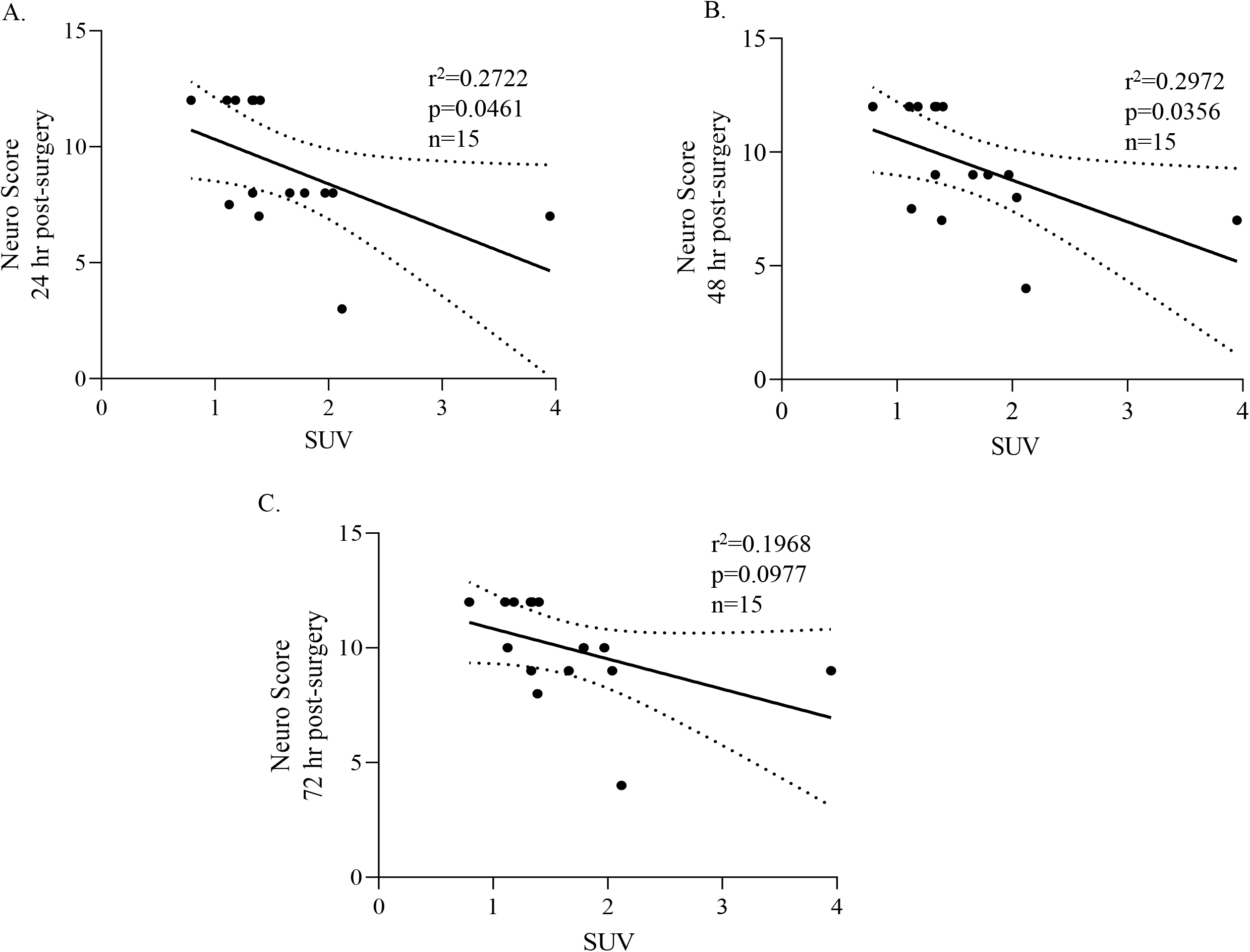
The increase in brain [^18^F]FDG uptake correlates with neurological injury induced by CA. Linear regression correlations between the whole brain SUV at 72 hour post CA and neuro scores at different time points post CA were plotted. Correlation of increased brain [^18^F]FDG uptake with lowered neurological scores was found for all the plots. A) & B) The statistically significance was achieved at 24 and 48 h post-CA. C) No statistical significance was observed at 72 h post-CA despite a trend is evident.

Figure 4C shows the representative coronal, transverse, and sagittal sections (left, middle, and right) of a CA mouse brain that were over laid with the 3D brain atlas. The SUV of all 14 regions are presented in Figure 4A. Overall, the lowest uptake was seen in the cortex with median SUVs of 1.03 for shams and 1.56 for CA animals. The medulla region had the highest [^18^F]FDG uptake with median SUV values at 1.45 for sham mice and 2.38 for CA mice. Compared to shams, all 14 brain regions of CA animals showed increased SUV. Statistically significant differences in SUV uptake were noted in 8 out of 14 regions (p<0.05) including the midbrain, pons, medulla, cerebellum, thalamus, corpus callosum, ventricles, and olfactory (Fig 4A). The midbrain and pons had the largest differences (43.2% & 43.1% respectively) among all 14 regions while the pallidum had the smallest difference (34.1%) (Figure 4B). The relative whole brain and regional [^18^F]FDG uptake using each individual liver as a reference region is shown in Figure 5. Similar to the SUV, the relative global cerebral [^18^F]FDG uptake was significantly higher in CA mice than the shams (p<0.05) (Figure 5A), while 3 (midbrain, pons, and medulla,) out of 14 regions showed significantly elevated[^18^F]FDG uptake (p<0.05) (Figure 5B).

**Figure 4.**
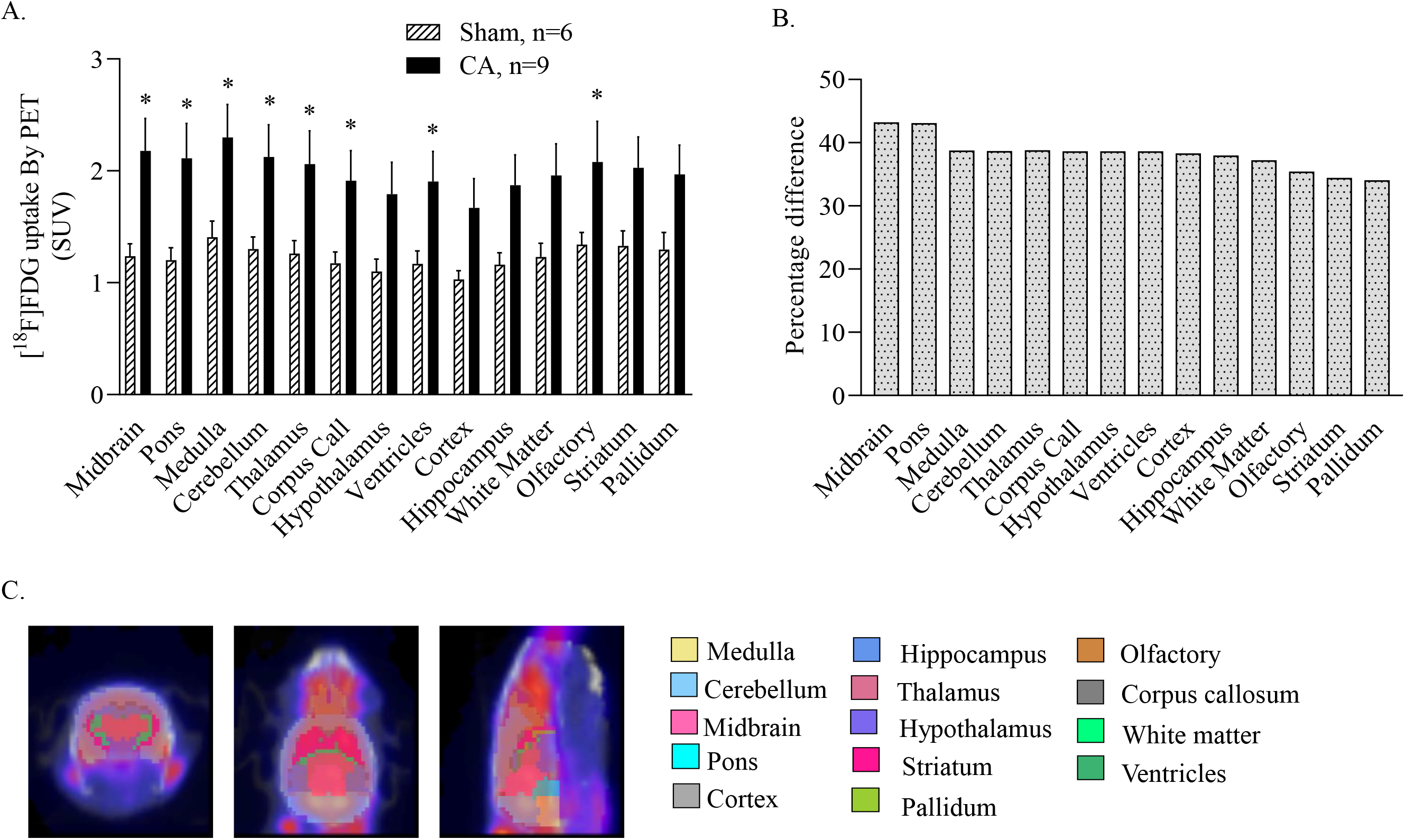
Regional analyses of the brain [^18^F]FDG uptakes. A) SUV of [^18^F]FDG uptake for 14 functional brain regions segmented using a 3D brain atlas. Eight of 14 regions showed a significant increase in [^18^F]FDG uptake in CA mice compared to sham mice. B) Percentage differences of the mean SUV between CA and sham animals. C) Representative coronal, transverse and sagittal sections of brain images overlaid with the 3D brain atlas.

**Figure 5.**
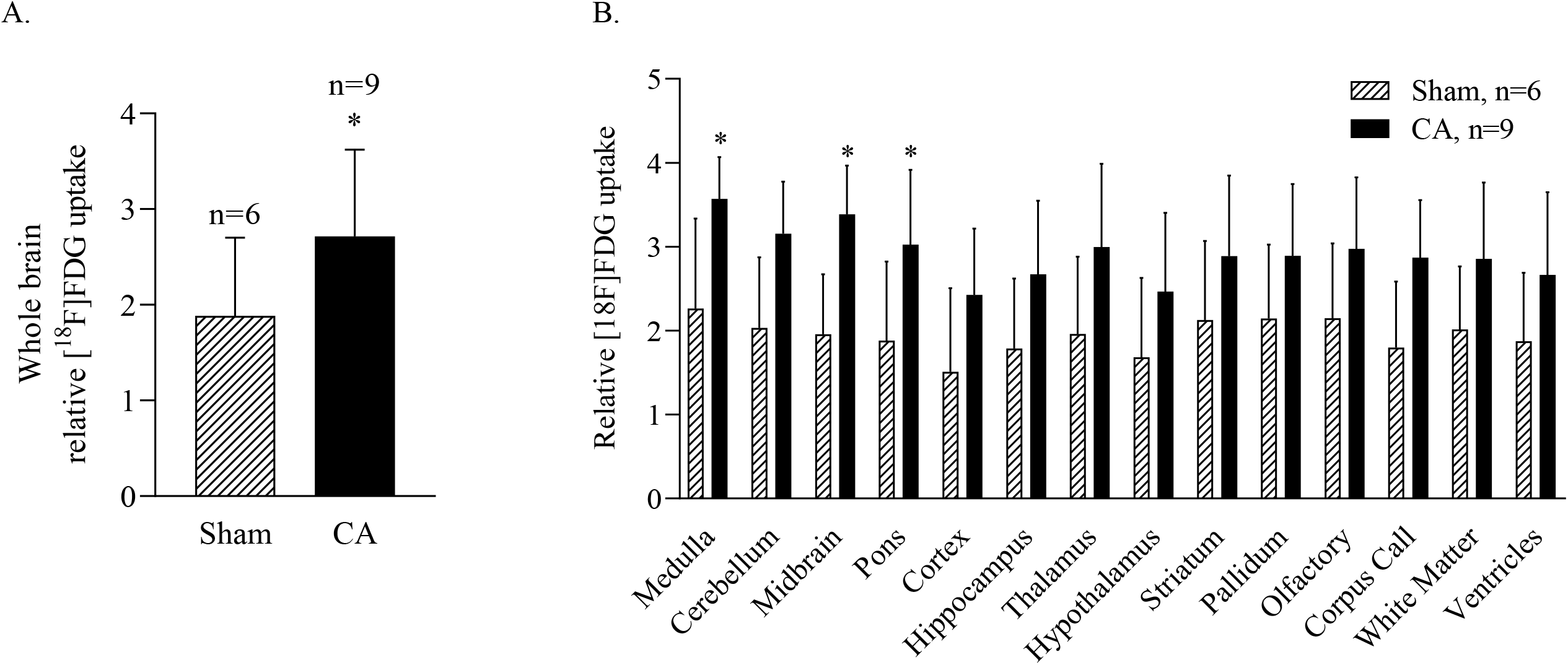
Relative brain [^18^F]FDG uptakes using liver as reference. A) Whole brain relative [^18^F]FDG uptake. B) Regional relative [^18^F]FDG uptakes.

## Discussion

According to a 2015 American Heart Association report, only 8.3% of those who survive cardiac arrest return to their former quality of life^21^. The majority of cardiac survivors sustain certain degrees of neurological injuries ranging from mild cognitive defects, impaired consciousness, to more serious conditions such as remaining in a persistent vegetation state. To date, the only therapies for this injury is conservative intensive care unit care and therapeutic hypothermia, yet outcomes remaining highly variable.

The brain consumes 25% of total body glucose as its primary source of energy^22,23^, even though it only makes up 2% of total body weight. Therefore, quantification of glucose metabolism may be a good indicator of overall brain function. Using the clinically available PET tracer, [^18^F]FDG, we examined changes in brain glucose metabolism in response to post-CA resuscitation. In our study, surviving post-CA mice had evidence of neurological injury as reflected by their neurological scores (Fig 1C). In contrast to our expectations of observing decreased brain metabolism in post-CA mice, we discovered significant increases in [^18^F]FDG uptake in the brain of post-CA animals at 72 hours.

The etiology of the increased metabolic demand in neurologically impaired animals at 72 hours post-CA is uncertain. Possible explanations for these observations could include increased neuro excitotoxicity following neuronal ischemia-reperfusion injury. Another explanation could be the increased seizure activity which has been reported in other rodent models of cardia arrest^24^. Although we did not directly observe seizure in our post-CA animals, we did not systematically check our mice for seizure activity. Seizure activity is thus a distinct possibility. Alternatively, post-CA inflammation in the brain may be responsible for the increased [^18^F]FDG uptake. Clinically, [^18^F]FDG is used as a biomarker for inflammatory diseases^25^. It is possible that infiltrating macrophages or the intrinsic activation of microglia resulted in the observed changes. Finally, depression of mitochondrial function with subsequent increases in glycolysis potentially results in increased brain glucose uptake. In 2010, Xu et al. reported a suppressed brainstem mitochondrial respiratory rate four days following cardiac arrest and resuscitation in aged rats^26^. Interestingly, we too found that the regions consisting of brainstem (midbrain, pons, and medulla) presented the largest and statistically significant increase of relative glucose uptake using liver as reference. It is thus interesting to speculate whether mitochondrial respiration is suppressed by CA induced brain injury in our animal model, especially in the brainstem region. Our results in this study could therefore reflect an increase in glucose demand due to a switch of glucose metabolism from oxidative respiration to glycolysis, similar to the Warburg effect in caner.

Currently, prognosis of cardiac arrest patient outcome is largely dictated by the distribution and degree of cerebral ischemic injury^27^. Much of the research focuses primarily on changes in cerebral blood flow or neuroanatomy. Brain function measurement for patients suffered CA is not clinically available. Here, we established a clear correlation between cerebral glucose uptake and CA induced neurological injury using [^18^F]FDG-PET imaging. When assessing differences of [^18^F]FDG uptake within specific brain regions, statistically significant elevations were found in 8 out of 14 regions analyzed.

Clinically, cerebellum, thalamus, corpus striatum, and hippocampus are known to be vulnerable to CA induced injuries^28^. In particular, imaging studies using diffusion-weighted magnetic resonance (MRI) consistently report reduced diffusion within the *cornu Ammon* neurons of the hippocampus, cerebellar Purkinje cells, and cerebral cortex ^29–32^. The finding in this study correlates with clinical outcomes in human patients for CA.

In two previous pre-clinical studies using piglet and rat models of cardiac arrest, lowered cerebral uptake of [^18^F]FDG 2 to 48 hours after CA were observed^5,6^. In contrast, our study demonstrated statistically significant 1.43 and 2.29 fold increases of whole brain [^18^F]FDG uptake in CA mice compared to sham and control mice respectively (median SUV 1.79 vs 1.25and 1.79 vs 0.78) 72 hours after CA. Our study differed from these previous studies in several important aspects which may explain the differences in our findings. First, our study performed neurological assessments of sham and post-CA animas which the prior studies lacked. We found that our observed changes in brain metabolism correlated with the severity of neurological outcomes of the mice. Second, our study used a longer cardiac arrest duration (12 min vs 6-8 min), possibly resulting in more severe injury. Third our [^18^F]FDG-PET imaging was done much later following cardiac arrest (72 hours vs 2-48 hours) and possibly reflects progressing late brain injury which often does not become apparent until days following cardiac arrest. Importantly, Putzu et al. showed increased relative brain [^18^F]FDG uptake in several rat brain regions (midbrain, pons, medulla, and cerebellum), which are similar to our regional analyses, despite a conclusion of lowered cerebral [^18^F]FDG uptake^5^.

### Study Limitations

There are several limitations of this study. First, the animal numbers in the study were relatively small with only total 25 mice used. PET-CT studies are technically challenging and expensive limiting the number of mice that we could analyze practically. Despite the relative small number of mice used in the study, we found significant changes in glucose utilization in response to CA, indicating the importance of further study. Secondly, we only analyzed brain glucose utilization at only one time point; 72 hours. We chose 72 hours for our study because prior studies in other rodent models had already looked at brain glucose uptake at 24 and 48 hours and because post-CA brain injury is clearly present on CT in both animals and patients at 72 hours post CA. Since our findings are in contrast to previous published findings, future studies examining time course changes in brain glucose utilization are warranted. Finally our study does not demonstrate the mechanism or anatomical nature of the injury observed in our mice behaviors and changes in the CT imaging which will be addressed in future.

## Conclusions

Asystolic cardiac arrest induced increases in global brain glucose uptake 72 hours following CA resuscitation that were associated with neurological injury. Increased brain glucose uptake was particularly significant in the brainstem, although widespread throughout the brain. The etiology of these changes is uncertain and requires further study.

## Acknowledgements

The authors acknowledge the assistance from the Integrative Small Animal Imaging Research Resources (ISAIRR) of the University of Chicago supported in part by the NIH grant P30 CA14500. The authors thank Nathanial Holderman and Andrew McVea for technical support.

---

**Supplemental Figure 1.**
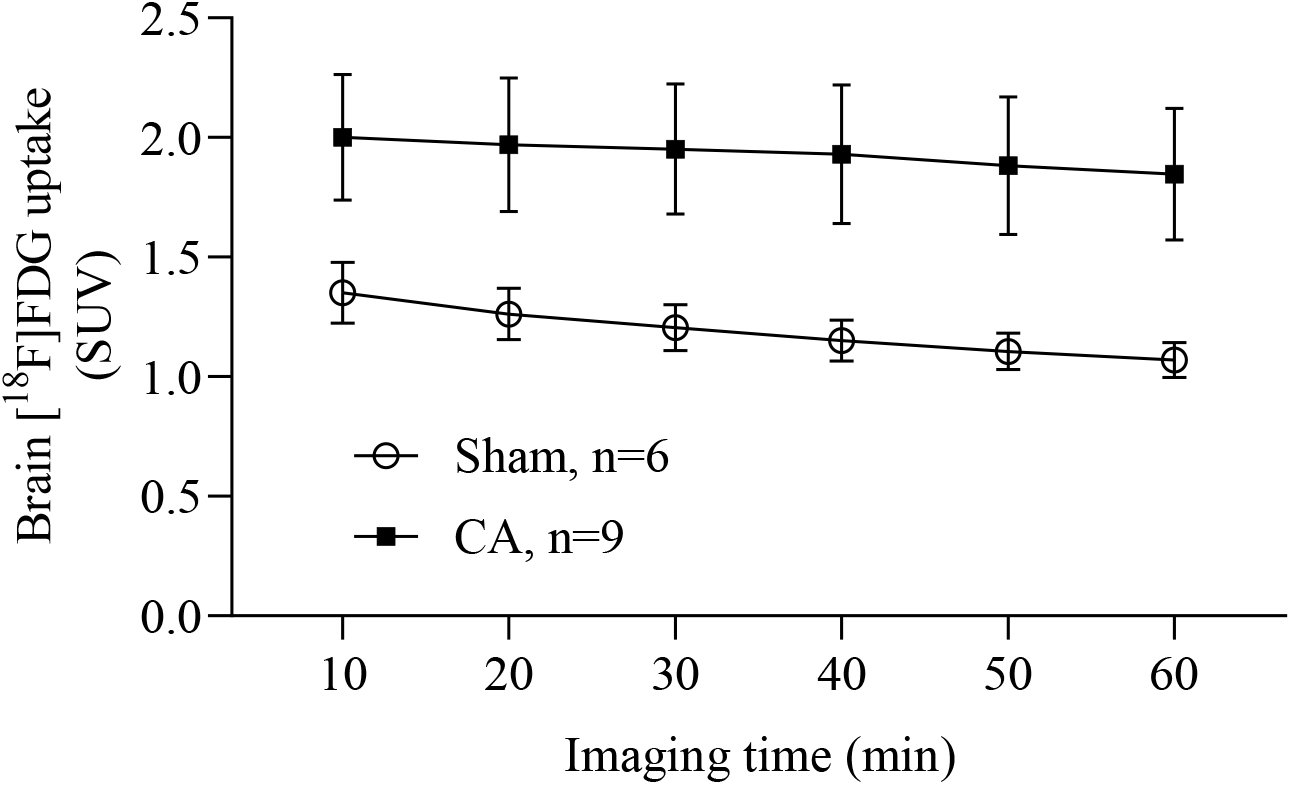
Average time active curve of whole brain [^18^F]FDG uptake

**Supplemental Figure 2.**
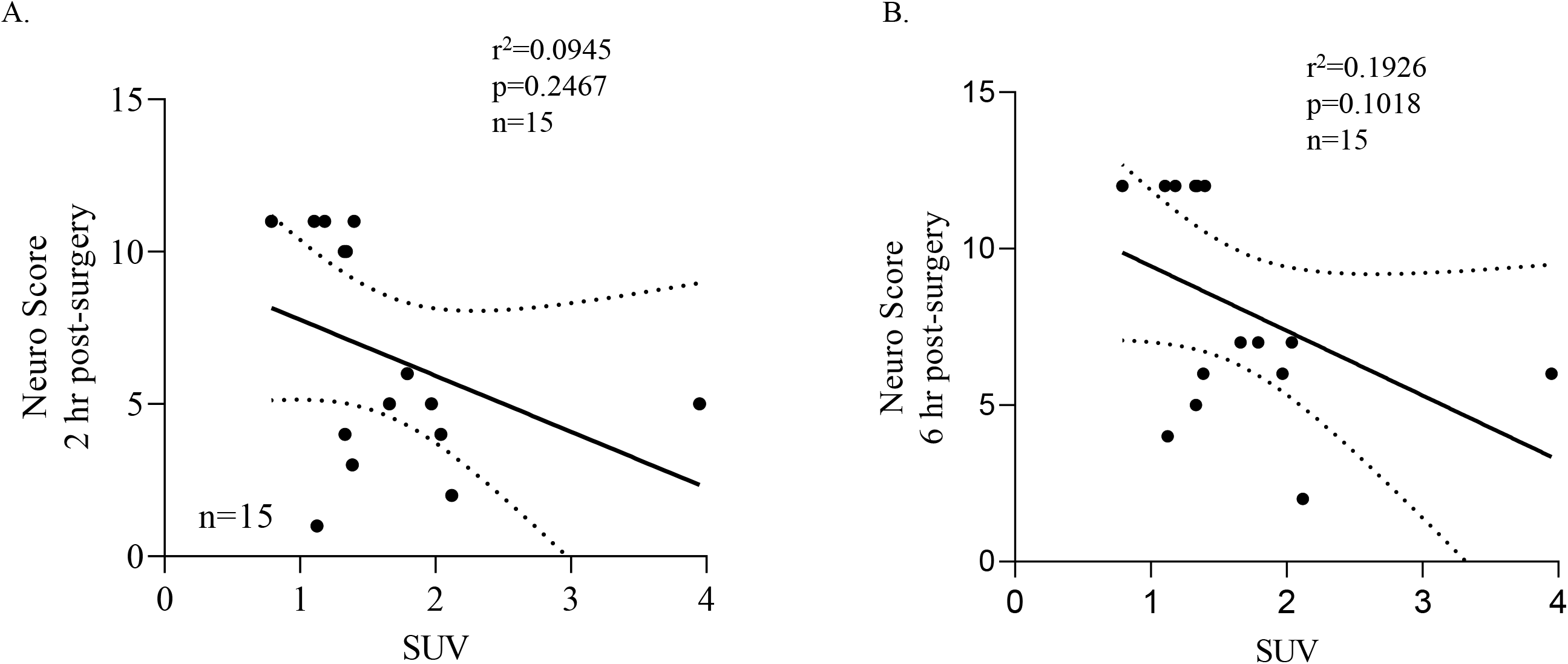
Linear regression correlations of increased brain [^18^F]FDG uptake with lowered neurological scores was observed at 2 h(A) and 6 h (B) post CA, but was not statistically significant.

